# Amplitude discrimination is predictably affected by echo frequency filtering in wideband echolocating bats

**DOI:** 10.1101/2021.10.04.463049

**Authors:** Amaro Tuninetti, Andrea Megela Simmons, James A. Simmons

**Affiliations:** Department of Cognitive, Linguistic, & Psychological Sciences, Brown University, Providence RI 02912; Carney Institute for Brain Science, Brown University, Providence RI 02912; Department of Neuroscience, Brown University, Providence RI 02912

## Abstract

Big brown bats emit wideband frequency modulated (FM) ultrasonic pulses for echolocation. They perceive target range from echo delay and target size from echo amplitude. Their sounds contain two prominent down-sweeping harmonic sweeps (FM1, ∼55-22 kHz; FM2, ∼100-55 kHz), which are affected differently by propagation out to the target and back to the bat. FM2 is attenuated more than FM1 during propagation. Bats anchor target ranging asymmetrically on the low frequencies in FM1, while FM2 only contributes if FM1 is present as well. These experiments tested whether the bat’s ability to discriminate target size from the amplitude of echoes is affected by selectively attenuating upper or lower frequencies. Bats were trained to perform an echo amplitude discrimination task with virtual echo targets 83 cm away. While echo delay was held constant and echo amplitude was varied to estimate threshold, either lower FM1 frequencies or higher FM2 frequencies were attenuated. The results parallel effects seen in echo delay experiments; bats’ performance was significantly poorer when the lower frequencies in echoes were attenuated, compared to higher frequencies. The bat’s ability to distinguish between virtual targets at the same simulated range from echoes arriving at the same delay indicates a high level of focused attention for perceptual isolation of one and suppression of the other.

## I. INTRODUCTION

Big brown bats (*Eptesicus fuscus*) are North American insectivores that navigate and forage using echolocation to build a perceptual image of the surfaces and environment around them. Their echolocation calls are wideband, frequency-modulated (FM) pulses ranging in duration from 0.6 ms to 20 ms, and sweeping downwards in frequency from around 100 kHz to around 22 kHz (Griffin, 1958; Simmons and Stein, 1980; Surlykke and Moss, 2000). The downward FM sweep consists of two to three harmonics: the second harmonic (FM2) sweeps downward from ∼100 kHz to ∼55 kHz, and the first harmonic (FM1) sweeps downward from ∼55 kHz to ∼22 kHz. A segment of the third harmonic (FM3) often is present, too, sweeping downward from ∼110 kHz to ∼80 kHz, but it is weaker. The frequencies above 50 kHz (i.e. above FM1) become very quickly attenuated by the atmosphere during propagation from the bat to a target and back (Griffin, 1958; Lawrence and Simmons, 1982; Stilz and Schnitzler, 2012). Beyond distances of a few meters, echoes contain most of their energy in FM1. Several experiments have demonstrated the bat’s emphasis of FM1 for perceiving target range from echo delay (see below; Moss and Schnitzler, 1989; Bates and Simmons, 2010; Bates *et al*., 2011; Stamper *et al*., 2009), The echo stimuli used in these previous experiments were varied in delay, providing a time offset between the positive and negative virtual targets that likely helped the bat to isolate the desired object for perception. In the new experiments reported here, we explore the relative roles of FM1 and FM2 in mediating the bat’s ability to discriminate the amplitude of echoes for target size. Here, the bats were presented with virtual targets 83 cm away, from echoes that arrived at the same delay of 4.8 ms. The simultaneity of echo arrival from both positive and negative stimuli adds the challenge of clutter suppression because each set of echoes arriving at the same delay could interfere with perception of the other set.

Big brown bats perceive the egocentric distance of acoustically-reflecting surfaces from echo delay with very high accuracy (Simmons, 1973; Moss and Schnitzler, 1989; Simmons *et al*., 1990). The wideband structure of their echolocation pulses, which cover a large number of frequencies in a short period of time, allows for more accurate distance measurements than do narrowband signals (Simmons, 1973; Simmons *et al*., 1975, 2004; Simmons and Stein, 1980; Boonman and Ostwald, 2007; Denny, 2007; Jones, 2008; Ming *et al*., 2021). The bats’ accuracy in perceiving a target’s egocentric distance (perceived from the time delay between outgoing pulses and returning echoes) has been investigated using psychophysical tasks (Moss and Schnitzler, 1989; Simmons *et al*., 2004; Stamper *et al*., 2009; Bates and Simmons, 2010; Bates *et al*., 2011) in which bats are trained to detect and discriminate virtual echoes based on pulse-echo delay, or to determine whether the pulse-echo delay changes from pulse-to-pulse (i.e. the target echo’s delay ‘jitters’ back and forth). Once bats have been trained to reliably discriminate two echoes (a target, rewarded, echo from a non-target echo), the echoes can be modified and the change in discrimination performance (if any) measured. With this paradigm, researchers can make assumptions as to how the bats’ perception changes as a function of the acoustic content of incoming echoes. These studies quantified the delay resolution of FM echolocating bats (∼10 ns) and revealed an ecologically relevant asymmetry in the perceptual role of higher and lower frequencies in determining pulse-echo delay.

Simmons et al. (2004) trained big brown bats to discriminate echoes with a set pulse-echo delay from echoes whose temporal delay jittered back and forth on subsequent echo presentations. When echoes were unfiltered, the bats could discriminate a non-jittering echo from a jittering echo when the jitter delay was at least 10 ns – equivalent to a change in distance of 0.0035 mm. When echoes were increasingly highpass filtered (from 15 – 35 kHz, in 5 kHz increments), discrimination thresholds steadily increased eightfold to 80 ns. Bates and Simmons (2010) replicated this effect in a non-jitter delay discrimination task. Big brown bats were instead trained to discriminate two simultaneous echoes separated by 800 µs (corresponding to 14 cm of physical distance between targets). As more of the lowest frequencies in the target echo were progressively highpass filtered, the bats’ discrimination performance progressively worsened. When echoes were filtered to only include frequencies between 66-90 kHz (i.e. FM1 was fully removed), the bats’ performance decreased below 50%, indicating that they switched to responding to the unfiltered echo as the target echo – despite it not being at the echo delay to which they had been trained to respond. These results suggest that without the lowest band of frequencies in an echo, the bat does not perceive the stimulus as an echo, and is thus unable to calculate pulse-echo delay.

In contrast to FM1, the higher frequencies of FM2 (∼100-55 kHz) are neither necessary nor sufficient for successful perception of echoes; that is, bats can still perform discrimination tasks if FM2 is absent, but not if echoes consist of *only* FM2 (Moss and Schnitzler, 1989; Stamper et al., 2009). Moss and Schnitzler (1989) trained big brown bats to discriminate between an echo with a constant delay and a jitter-delay echo, where the jitter delay was between 0.4 – 4.8 µs. When echoes were highpass filtered at 40 kHz, requiring the bats to discriminate echo delay using primarily FM2, the bats “failed to perform” and “refused to make a choice” (p. 389). Thus, performance was dependent on the presence of FM1. Bates and Simmons (2010) found similar results – the bats’ performance in a discrimination task did not worsen when echoes were filtered to only include FM1.

It is not the case, however, that the frequencies contained in FM2 are not perceptually informative to the bat. Stamper et al. (2009) found that, when FM2 was split from FM1 and delayed in time (relative to FM1), the bats made more errors when a non-target echo coincided with the delay of that split-harmonic echo. These results suggest that the upper frequencies of FM2 influence the bat’s perception of echo delay (or distance from the bat) if they are present, but the bat is also able to perceive distance using only the lower frequencies of FM1, if necessary. Additionally, delay-accuracy for split-harmonic echoes was overall worse than for harmonically-aligned echoes, suggested that temporal alignment of echo frequencies is required for highly accurate perception of echo delay.

Bates et al. (2011) ran a series of experiments showing that the upper frequencies of FM2 affect echo perception in a more graded manner than the frequencies of FM1, which completely disrupt the bat’s perception when absent. When FM2 of a non-target echo was not removed or delayed, but attenuated (i.e. weakened), delay discrimination performance approached 100%, suggesting that their temporal perception of the non-target had become defocused as a result of the attenuation of its higher frequencies. These results, along with those described above, outline a comprehensive perceptual clutter rejection mechanism which allows bats to perceive the object ensonified by the center of their echolocation beam with high temporal acuity, while simultaneously temporally defocusing more peripheral echoes (the more peripheral, the more defocused) so that these incoming peripheral echoes do not mask the bat’s highly accurate delay percept of the center of the beam (Bates et al., 2011).

In the current experiment, we aimed to extend these previous results in a different perceptual discrimination context. Rather than using an echo-delay discrimination task, we tasked bats to discriminate virtual targets on the basis of amplitude, which corresponds to the perceived size of an ensonified object (Simmons and Vernon, 1971).

## II. METHODS

### A. Animals

Five adult big brown bats (named F., G., J., K., and M.; four females and one male) were trained for this experiment. They were wild-caught from barns or attics in Rhode Island under a state scientific collecting permit. Because they were wild caught, their ages are unknown beyond one year. Bats were housed in groups of 2-3 individuals in a temperature- and humidity-controlled colony room (22-25° C, 40-60% humidity) on a 12:12 reversed dark:light cycle. Individuals were identified by scannable microchips implanted subcutaneously in their upper backs over one month before the experiment began. They had unlimited access to vitamin-enriched water and received their daily food allotment (live mealworms, *Tenebrio larvae*) during experiments as rewards for correct performance. Bats were not food-deprived throughout the duration of the experiment and were maintained at healthy weights between 15.0 and 18.0 g. All procedures were approved by the Brown University Institutional Animal Care and Use Committee and are consistent with federal guidelines.

### B. Virtual target presentation system

Bats were trained to complete a two-alternative forced-choice (2AFC) task which required them to choose the stronger (higher amplitude) of two ultrasonic echoes, or virtual targets. The task took place in an 8.3 m × 4.3 m × 2.7 m room lined with sound-absorbent foam (SONEX) on the ceiling and walls and artificial athletic turf on the floor to attenuate unwanted echoes. The 2AFC platform was located on the room’s midline, 5.4 m from the back of the room (the direction the platform faced), and at a height of 1.2 m from the floor. There was 1.2 m of empty space on either side of the platform, and 4.0 m of empty space to the front of the platform, so as to avoid extraneous room echoes reaching the bats at similar time delays as the experimental stimuli. The room was illuminated with dim, long-wavelength red light to allow for bat handling and video monitoring by the experimenters.

Each bat was trained to sit at the base of an elevated Y-platform and broadcast its echolocation calls towards the end of the platform (Fig. 1). At each end of the platform’s two arms was an ultrasonic microphone (Knowles Electronics FG-3329), separated from the other by 11 cm and 29° (relative to the point at which the bat crawls onto the platform). These microphones recorded the bat’s echolocation calls and immediately delivered them back to the bat as virtual echoes from two ultrasonic speakers (Tucker-Davis ES1, 3.8 cm diameter), mounted 1.4 m from the edge of the platform (Fig. 1). The two speakers were placed 86 cm and 35° apart, with each speaker aimed directly at its corresponding platform arm. Ultrasonic calls recorded by the left platform microphone were routed to the left speaker, and vice versa. Each speaker was mounted 1.4 m from the edge of the platform to create a time delay between the bat emitting echolocation calls and the bat receiving the corresponding delivered echoes. This distance, combined with the distance that the echolocation calls had to travel to reach the platform microphones, resulted in a total pulse-echo delay of approximately 4.84 ms, corresponding to a pair of virtual targets presented at a distance of ∼83 cm from the point at which the bat walks onto the platform (Fig 1).

**FIG 1.**
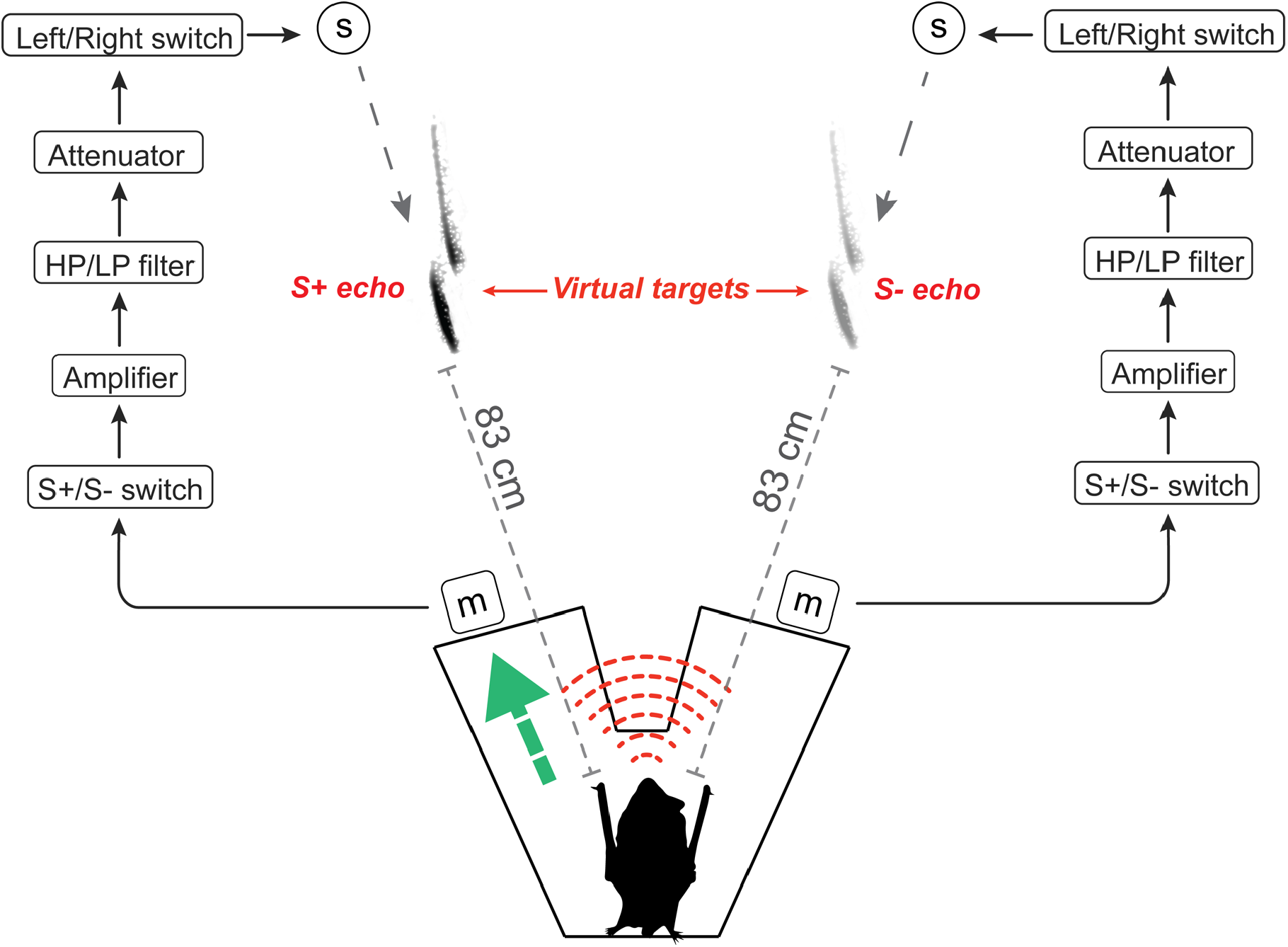
Diagram of experimental setup. A bat sitting on a Y-platform emits ultrasonic echolocation calls (red dashed lines; color online) which are picked up by two microphones (m) and simultaneously emitted from two ultrasonic loudspeakers (s) mounted at a distance of 1.4 m from the platform, to create two virtual targets 83 cm in front of the bat (S+ and S-, shown as call spectrograms). Labeled boxes indicate signal processing equipment used to amplify and filter each of the audio channels. Five bats were rewarded for walking (green arrow; color online) in the direction of the stronger of two echoes (S+, denoted by a darker spectrogram). Bats were not rewarded for walking in the direction of the weaker echo (S-, denoted by a weaker spectrogram). The direction (left or right) of the S+ and S- echoes was counterbalanced across trials according to a pseudorandomized schedule (Gellermann, 1933).

The emitted calls recorded by each platform microphone were highpass filtered at 10 kHz (ThorLabs EF121 HP filter) to remove background noise, routed to a microphone preamplifier (RME 4-channel Quadmic preamplifier), and then into a custom-built switchbox (Fig. 1, “S+/S- switch”) which designated each of the two audio channels carrying the bat’s calls (left and right platform microphones) as either the positive stimulus (S+) or the negative stimulus (S-). The switchbox thus determined the direction of the S+ and S- stimuli for each trial (if the switch was positioned to the *right*, the *right* speaker emitted S+ and the *left* speaker emitted S-, and vice versa if the switch was positioned to the left). After routing through the switchbox, both stimulus channels were then further amplified, filtered, and attenuated, with parameters varying by experimental condition (see Experimental stimuli section). Amplification of each channel was accomplished with two preamplifier units (FMR Audio, RNP8380), filtering of either the S+ or the S- channel was accomplished using two consecutive analogue filters (Rockland Model 852 Dual hi/lo filter, combined 96 dB/octave), and attenuation of each channel was accomplished with two attenuators (Tucker-Davis Technologies, PA5 programmable attenuator). All amplification and filtering equipment was located outside the experimental room. After S+ and S- channels were appropriately filtered and attenuated, they were again routed through the custom-built switchbox (Fig. 1, “Left/Right switch”) to re-designate the S+/S- channels as Left/Right audio channels for emission through their corresponding speaker. During training and data collection, the direction of S+ and S- was pseudorandomly varied from trial-to-trial according to a Gellermann (1933) schedule. After S+ and S- are assigned to a Left/Right speaker, both stimuli were routed to a two-channel speaker driver (Tucker-Davis Technologies, ED1 electrostatic driver) and then emitted through their corresponding speaker in front of the platform. Each of the four audio channels described here (left and right platform microphones, left and right speakers) were also recorded on an audio recorder (Zoom F4, digitized at 192 kHz) to analyze the spectral content of calls emitted by the bat and the resulting echoes emitted by speakers.

The system was calibrated using a 2 ms-long, 2-harmonic FM sweep from 100-20 kHz (i.e. an artificial echolocation pulse) synthesized in Adobe Audition (2019) and generated at 2.0 V by a digital signal generator (Koolertron, JDS2600-60M). This signal was inserted into the system in lieu of actual bat calls, and the strength of the emitted echoes from each individual speaker was calculated by placing an ultrasonic microphone (Brüel & Kjær Model 4135 1/4-inch) in the center of the platform facing the speakers. A calibration signal generated from the signal generator at 2.0 V was comparable in amplitude to the strongest bat calls emitted during the experiment (measured by oscilloscope during pilot trials), and resulted in echoes of 78 dB SPL at the platform, indicating that echoes reaching the platform were well above the hearing threshold of big brown bats (Koay et al., 1997). Calibration measurements were run with only one stimulus (S+ or S-) present in order to measure the amplitude of each stimulus individually, rather than the amplitude of both stimuli arriving at the platform at the same time. This calibration method was also used to measure the decrease in echo amplitude after echoes were high- or lowpass filtered, in order to compensate for the reduced acoustic energy present in echoes after filtering.

### C. Training and data collection

Two experimenters were present on each day of training and data collection, and trials were run using a double-blind procedure. Experimenter 1 handled the bat on each trial and was blind to the experimental sequence and the correct choice for all trials. Experimenter 2 was positioned behind Experimenter 1, separated by an opaque felt screen, and monitored the bat’s response via a ceiling-mounted black and white CCD video camera (DSP 15-CB22 1/3” sensor B/W camera), which provided a live bird’s-eye view of the platform to a video monitor (Blackmagic Video Assist). Experimenter 2 controlled the left/right position of the positive (S+) and the negative (S-) stimuli according to a prearranged pseudorandomized sequence (Gellermann, 1933) and verbally informed Experimenter 1 if the bat’s response was correct or incorrect after each trial.

Bats were trained 5 days a week to walk in the direction of S+. On each trial, the bat was rewarded with a piece of mealworm for walking down the arm of the platform which corresponded to the speaker delivering the S+ echo. If the bat walked down the arm corresponding to S-, a broadband ‘shh’ sound was made by Experimenter 1 to signal to the bat that it made an error, and the bat was held in the hand for a 5-sec interval before beginning the next trial. Training began with the S+ echo not attenuated (−0 dB on the corresponding Tucker-Davis attenuator) and the S- echo completely attenuated (−120 dB on the corresponding attenuator). Once a bat was able to correctly respond to (i.e. walk in the direction of) the S+ echo on 90% of trials in one day, the S- echo was introduced at the same overall pulse-echo delay as S+, but at −45 dB relative to S+. At this point, the bat had to distinguish between two echoes, both of which were present after each emitted echolocation call and at the same time delays (i.e., the same distances from the bat), but which differed in their amplitude.

Over the course of 8-14 weeks of training (the amount of training required varied per bat), the attenuation of the S- echo (relative to the S+ echo) was gradually reduced for each bat, leading to smaller amplitude differences between the two stimuli. Once a bat demonstrated its ability to discriminate S+ and S- at the test amplitudes (i.e. the bat walked in the correct direction on ≥75% of trials on a given day) for two consecutive days, the amplitude of S- was increased by 2-5 dB for that bat on the next day of training. The final two weeks of each bat’s training involved smaller S- amplitude changes (0.5-1.0 dB at a time) to avoid making the day-to-day changes in the task too difficult for the bat. This process continued until the attenuation of the S- echo relative to the S+ echo was small enough that the bat’s performance dropped *below* 75% for two consecutive days, indicating that the bat was no longer able to discriminate the two echoes well. For each individual bat, the (S+):(S-) amplitude difference which resulted in below 75% correct performance was deemed the amplitude discrimination limit (ADL) for that bat. While 75% correct performance is commonly used as the threshold for successful discrimination (Simmons, 1973), performance was variable and could not be held at exactly 75% from day-to-day. For this reason, attenuation differences were chosen that resulted in performance levels between 75% and 50% for each bat; this criterion avoided potential discrimination ceiling effects in the bats’ performance and is what we have defined as the bats’ ADL for the purposes of this experiment.

During training, each bat performed 10-50 trials per day (5-6 days per week). The number of trials a given bat performed on a given day was a function of the quantity of mealworms that bat was receiving as its daily food allotment and its experience with the task.

Once a bat reached its individual ADL during training, data collection began. There were five total experimental conditions, one of which was the “baseline” amplitude discrimination task – that bat’s performance at its measured ADL. The other four conditions covered all permutations of S+/S- filtering: S+ highpass filtered (HP), S- highpass filtered, S+ lowpass (LP), and S- lowpass filtered. In all conditions the amplitude difference between S+ and S- was maintained at the same level as in the baseline amplitude discrimination condition for each individual bat, such that the only difference between stimuli from condition to condition was the spectral filtering of either S+ or S-. The order of conditions was randomized for each bat. Bats participated in the four experimental conditions involving filtered stimuli every other day, with intervening days consisting of the basic amplitude discrimination task at (or near) that bat’s ADL. These intervening days of amplitude discrimination served to maintain the bats’ initial training of responding to the stronger of two echoes over the course of data collection. To maintain the bats’ robust amplitude discrimination training (acquired over 8-14 weeks of training), but also avoid frustrating them (which can occur with difficult discrimination tasks), intervening days were conducted with the S+/S- amplitude difference either at the bat’s ADL, or with S- further attenuated by 1-2 dB. These intervening days ensured that the bats completed the experimental discrimination tasks based on the amplitude difference between S+ and S-, and did not begin instead confounding any of the echo filtering conditions with food rewards.

Our goal was to collect 150 trials per bat in each condition, at a rate of 50 trials per day. This equates to three days of data collection per bat per condition, with one day of baseline amplitude discrimination between each day of data collection, to maintain their discrimination training. Unfortunately, data collection was halted by state-mandated stay-at-home orders precipitated by the COVID-19 pandemic, leading to an unequal number of trials across conditions. In the end, every condition contains data from at least three bats; in two conditions (S+ HP and S- LP) one of those bats did not reach 150 trials. Statistical power was calculated (G*Power 3.1, 2021) to confirm that all conditions consisted of enough trials to ensure adequate statistical power above 80% (with α = 0.05) for all statistical tests.

### D. Experimental stimuli

In the baseline amplitude discrimination condition, the stimuli that the bat received at the platform were spectrally and temporally identical to the echolocation calls recorded by the two platform microphones. The only aspect differentiating the S+ and S- echoes (i.e. the parameter the bats were trained to discriminate) was their amplitude, with non-target S- echoes being 3-6 dB weaker than target echoes (as determined by the individual bat’s ADL; see Results). In the four experimental conditions, either the S+ or S- echoes were also either highpass or lowpass filtered, while the amplitude of S- relative to S+ was maintained at the same level as in the baseline condition. The filtering of S+/S- was accomplished by routing either the S+ audio channel or the S- audio channel through two Rockland filters (Model 852 Dual hi/lo filter) set to the same settings, resulting in a 96 dB/octave attenuation beginning at the frequencies specified on the filter. For the two conditions requiring highpass filtering, the filter was set at 15 kHz above the lowest frequency in the bat’s echolocation calls; also known as the terminal frequency (TF) of the FM sweep. The TF of echolocation calls can vary between individual bats, so separate TFs were measured for each individual bat by visually inspecting the recorded spectrograms in Adobe Audition (2019). The mean TF for each bat was calculated by averaging the TF of all calls emitted in a single trial during that bat’s first day in the baseline amplitude discrimination condition. For the two conditions requiring lowpass filtering, the filter was set at 70 kHz for all bats. This frequency cutoff was chosen for all bats because the upper frequencies of big brown bat calls do not differ between individuals as drastically as the lowest frequencies; the upper frequencies extend into the call’s second or third harmonic and decrease in attenuation gradually, rather than abruptly ending as they do at the TF of the bat’s FM sweep. In contrast to the TF of a call, the presence or absence of these higher frequencies is more likely to be a factor of the strength of the emitted call, the distance the call has to travel, and the ensuing atmospheric attenuation, rather than a factor of any inter-individual differences in vocalization frequency. Moreover, the measurement of these highest frequencies depends largely on the sensitivity and sampling rate of the recording equipment used. Our recording sampling rate of 192 kHz made measurements above 96 kHz impossible, while the frequency responses of the ultrasonic microphones and speakers used begin to roll off above **∼**85 kHz. Ultimately, a 70 kHz lowpass limit was chosen because it resulted in a noticeable attenuation of a 10-15 kHz range at the upper limit of the emitted echoes, as judged by the spectrograms of the calibration signal and the bat calls emitted during trials. Fig. 2 provides examples of how the attenuation and filtering settings modulated S+ and S- in different conditions (example signals shown from one bat, M.).

**FIG 2.**
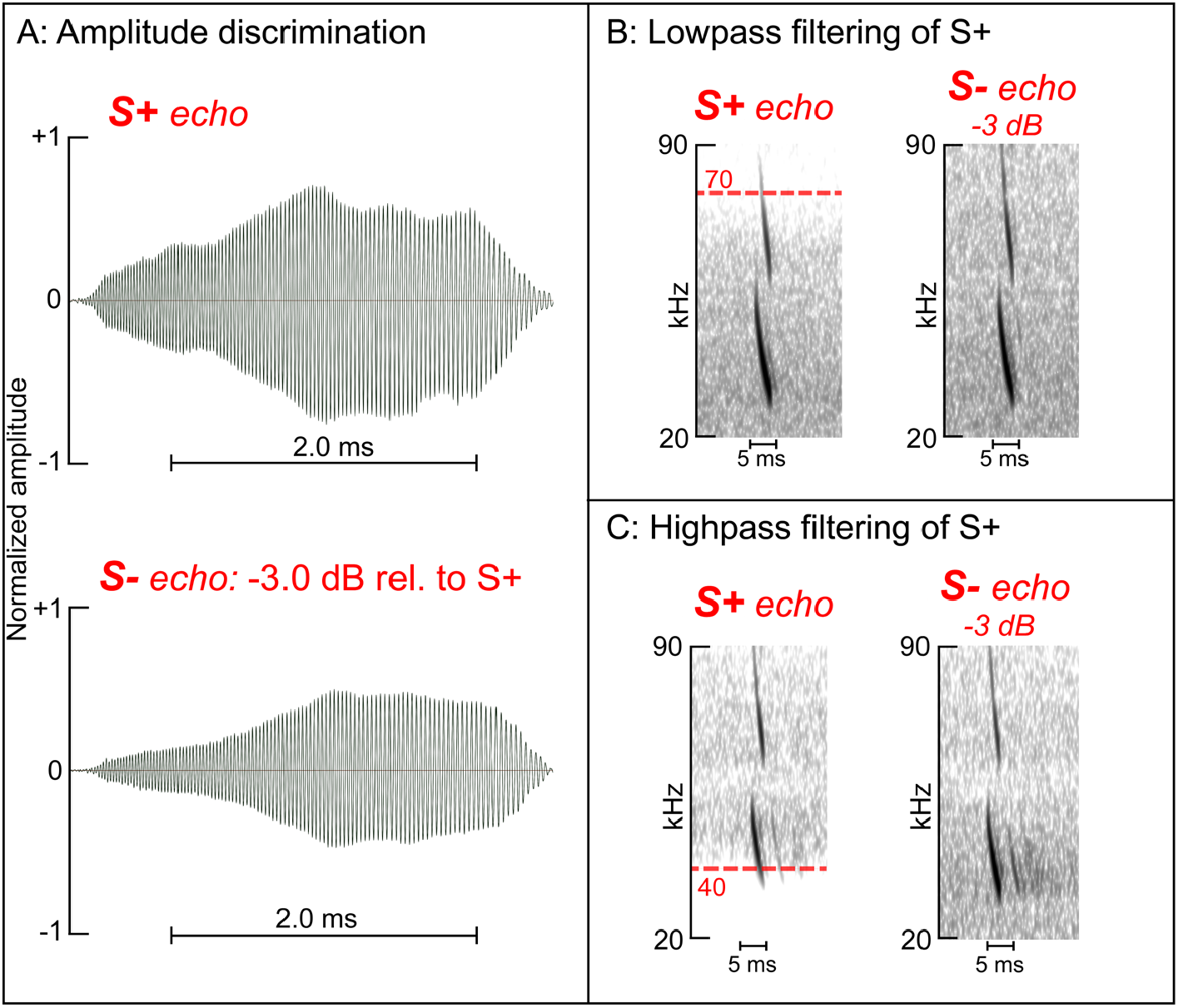
Example echolocation calls from one bat in three experimental conditions. (A) An example waveform of a single call from bat M. in the baseline amplitude discrimination condition. The same call from the bat was recorded by the two platform microphones and emitted by the two speakers to create two echo stimuli: S+ (top waveform) is unfiltered and unattenuated, (∼78 dB SPL at the bat), and S- (bottom waveform) is unfiltered but attenuated 3.0 dB relative to S+. Small differences in waveform shape are due to the call being recorded by two separate microphones, each at slightly different angles from the bat’s mouth at the time of call emission. (B) An example spectrogram of a single call (bat M.) in a lowpass (LP) filtering condition. LP filtering of the S+ echo at 70 kHz strongly attenuates the upper frequencies of the call’s second harmonic (left spectrogram). The other stimuli, S-, is unfiltered but is still attenuated 3.0 dB relative to S+, as in the baseline amplitude discrimination condition (see Methods). (C) A single call (bat M.) in a highpass (HP) filtering condition. HP filtering attenuates the lower end of frequencies (left spectrogram, HP at 40 kHz for this individual bat; see Methods and Results). The S- echo is unfiltered but remains attenuated by 3.0 dB relative to S+ in all conditions for this individual bat (see Methods, Results).

**FIG 3.**
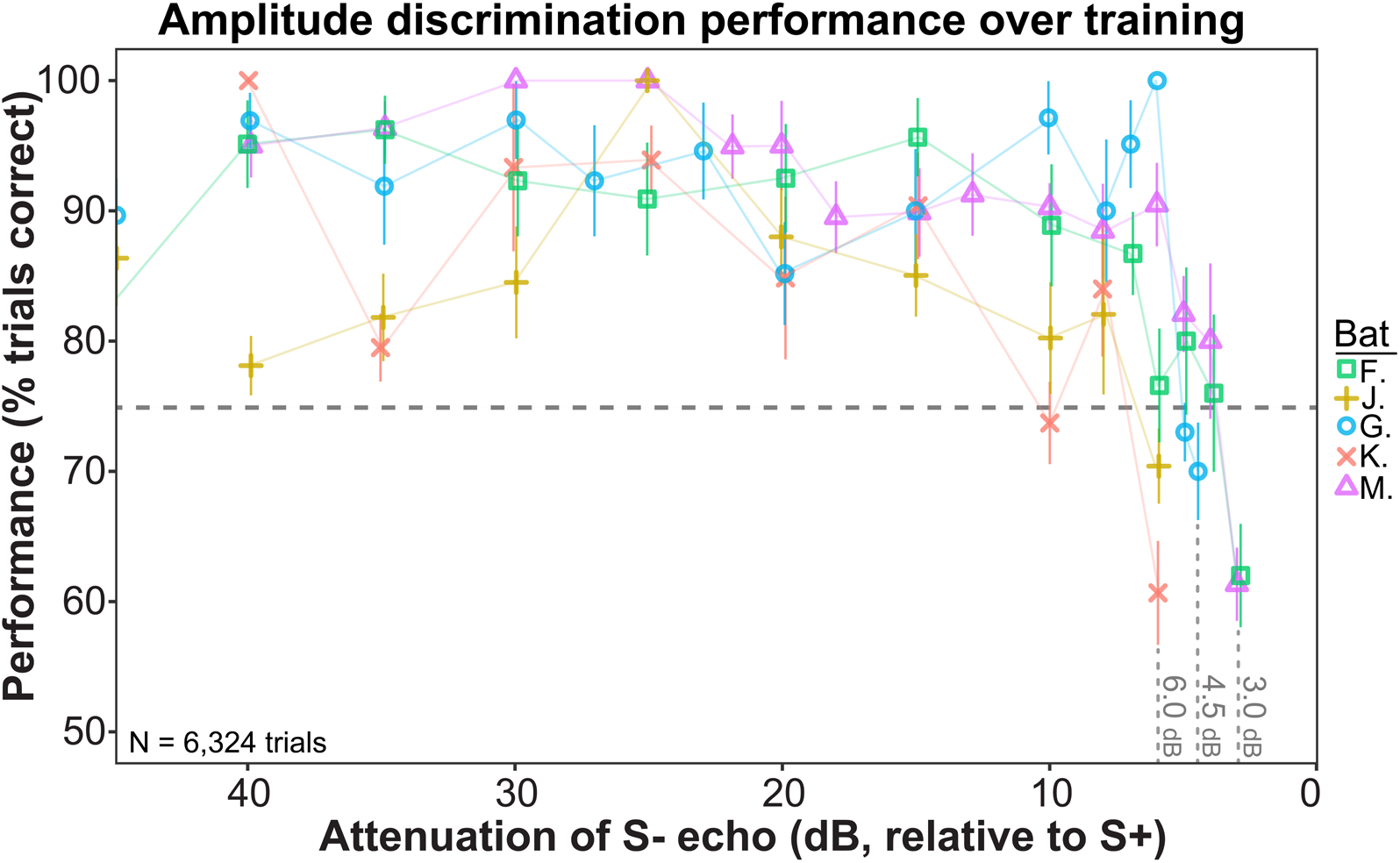
Bat performance on amplitude discrimination task over training period. Performance of each individual bat (different shapes, color available online) on the amplitude discrimination task over the course of training is plotted as a percentage of all trials performed at each amplitude difference level. Over 8-14 weeks of training, the attenuation of the S- echo (relative to the stronger S+ echo) was progressively decreased in steps of 0.5-5 dB, until the point at which the bat showed <75% discrimination performance for two consecutive days. Error bars indicate ±1 binomial standard deviation.

Because successful discrimination of echoes depended on small differences in amplitude, steps were taken to ensure that (S+):(S-) amplitude differences were maintained across conditions. Using the same calibration signal and microphone described above, we measured the decreases in stimulus amplitude caused by removing either upper or lower frequencies with analog filters. These slight decreases in amplitude as a result of filtering were then compensated for by increasing the amplitude of filtered echoes by the equivalent amount in each of the four conditions involving filtering.

### E. Data analysis and availability

Statistical tests were performed using RStudio (2018) and G*Power 3.1 (2019). All data are available in the Brown University data repository (https://doi.org/10.26300/c974-0k69). Performance data of each bat were input into RStudio (2018) and performance was calculated as the proportion of trials on which the bat responded correctly, per condition. Using a custom R script, mean performance (*p*) and binomial standard deviation (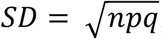, where *q* = 1 − *p*) were calculated for each condition, collapsing across bats. Two-tailed exact binomial tests were run to compare performance across conditions. In total, five binomial tests were run: one to compare mean performance in the baseline amplitude discrimination to chance (0.50), and four to compare mean performance in the baseline amplitude discrimination to mean performance in each of the filtering conditions. The statistical power of each exact binomial test, as a function of the number of trials collected and the proportions compared, was calculated with post hoc power analyses run in G*Power 3.1 (2020).

## III. RESULTS

### A. Amplitude discrimination limits and terminal frequencies

After 8-14 weeks of training (and decreasing amplitude differences between stimuli), none of the five bats were able to discriminate between S+ and S- echoes on the basis of amplitude (performance was <75% correct for two consecutive days). This S+:S- amplitude difference was defined as the amplitude discrimination limit (ADL) for each bat. Fig. 2 and Table 1 show the ADL reached by each bat: two bats reached an ADL of −3.0 dB, one bat −4.5 dB, and two bats −6.0 dB. Once a bat reached its ADL, the mean terminal frequency (TF) of its calls was calculated by averaging the TF of all calls emitted during one trial. These individual TF measurements affected the filtering for later highpass (HP) conditions; HP filtering was set to 15 kHz above the TF of each individual bat. Table 1 shows each bat’s ADL, calculated mean TF, and the filtering settings used for that bat.

**TABLE 1.** Amplitude discrimination limits (ADL) and filtering settings for individual bats. ADL describes the relative attenuation of the S- echo (relative to the S+ echo) at which that bat’s discrimination performance fell below 75% correct. HP filter frequency was set at 15 kHz above the terminal frequency (TF) of each individual bat’s echolocation calls, determined by averaging the TF of all calls emitted throughout one trial in the baseline condition. LP filter frequency was the same for all bats (70 kHz).

|  | Bat M. | Bat K. | Bat F. | Bat J. | Bat G. |
| --- | --- | --- | --- | --- | --- |
| **ADL (dB of S- attenuation)** | -3.0 | -6.0 | -3.0 | -6.0 | -4.5 |
| **Mean TF (kHz)** | 25.0 | 22.0 | 22.8 | 21.0 | 21.6 |
| **HP filter frequency (kHz)** | 40 | 37 | 37.8 | 37 | 37.6 |
| **LP filter frequency (kHz)** | 70 | 70 | 70 | 70 | 70 |

### B. Statistical power

Due to state-mandated COVID-19 restrictions, we were unable to collect the full number of planned trials from each bat across conditions. Table 2 outlines how many trials were collected in each condition from each bat, as well as the calculated statistical power achieved for each condition across all bats (calculated using G*power 3.1 software, assuming α = 0.05). Power for the baseline condition was calculated for a two-tailed exact binomial test comparing the baseline proportion of successes to chance (0.50). Statistical power for each filtering condition was calculated for a two-tailed exact binomial test comparing the proportion of successful trials in the experimental condition to the proportion of successes in the baseline discrimination condition. All calculations indicate that ensuing exact binomial tests have statistical power above 0.80 (*P* < 0.05).

**TABLE 2.** Number of trials achieved in each condition and corresponding statistical power. Post hoc power analyses were conducted to ensure that exact binomial tests had sufficient statistical power given the number of trials collected.

| Condition | Total # of trials | Power | # trials (Bat M.) | # trials (Bat K.) | # trials (Bat F.) | # trials (Bat J.) | # trials (Bat G.) |
| --- | --- | --- | --- | --- | --- | --- | --- |
| Baseline | 750 | 1.0, $p = 0.0445$ | 150 | 150 | 150 | 150 | 150 |
| S- HP | 600 | 0.99, $p = 0.0464$ | 150 | 0 | 150 | 150 | 150 |
| S+ HP | 436 | 1.0, $p = 0.0415$ | 150 | 136 | 150 | 0 | 0 |
| S- LP | 350 | 0.99, $p = 0.0457$ | 150 | 150 | 0 | 50 | 0 |
| S+ LP | 450 | 0.86, $p = 0.0448$ | 150 | 0 | 150 | 0 | 150 |

### C. Discrimination performance

Fig. 4 shows the performance of all bats in each of the discrimination conditions, as the percentage of successful trials in each condition. All comparisons between conditions were analyzed using two-tailed exact binomial tests. In the baseline amplitude discrimination condition, in which bats responded to the stronger of two echoes (S+) at their specified ADL (see Table 1), mean performance was 63.33% ± 1.76%, significantly higher than chance (*P <* 0.001). When S- was highpass filtered such that its lowest frequencies were removed, the bats’ performance increased significantly to 76.33% ± 1.74% (*P <* 0.001). When S- was unfiltered and S+ was instead highpass filtered, such that the lowest frequencies of S+ were removed, the bats’ performance decreased significantly from baseline performance to 32.34% ± 2.24% (*P <* 0.001), despite the fact that S+ had a higher amplitude than S- in this condition. When S+ was left unfiltered and S- was lowpass filtered to remove its highest frequencies, the bats’ performance again improved significantly relative to baseline performance, to 73.71% ± 2.35% (*P <* 0.001). When S- was unmodified and S+ was instead lowpass filtered, performance relative to baseline decreased significantly to 56.22% ± 2.34% (p = 0.002).

**FIG 4.**
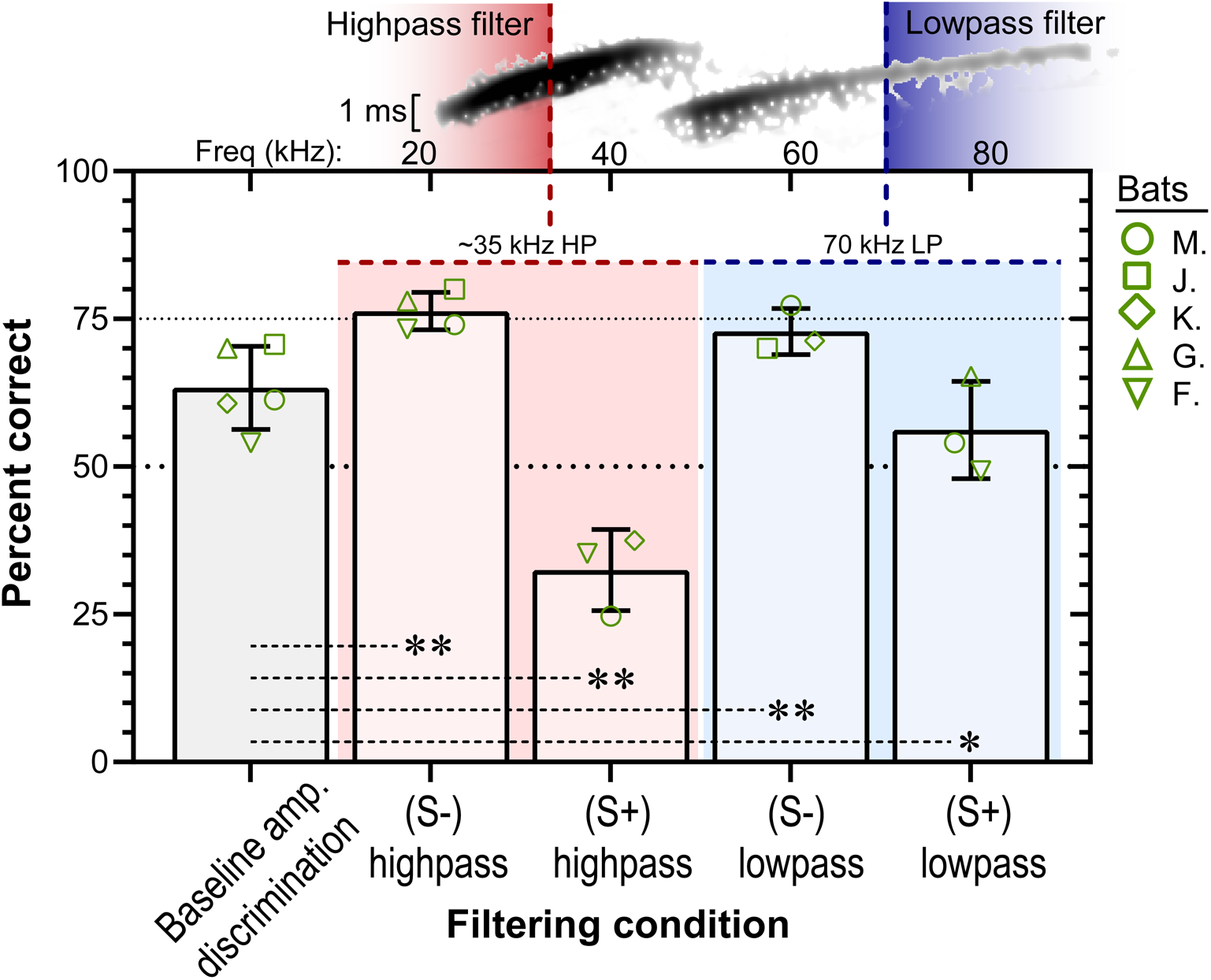
Performance of all bats in amplitude discrimination task. Baseline discrimination (grey bar) required responding to the stronger of two simultaneously-presented echoes. An example echo spectrogram (black) lies horizontally above the plot, with frequency in kHz plotted on the top axis and colored dashed lines indicating at what frequency echoes were filtered (highpass filter values vary slightly by bat; see Table 1). Highpass filtering (red shading; color online) removed the lower 15 kHz of echo frequencies, and lowpass filtering (blue shading; color online) removed the upper end of frequencies. Green symbols (color online) show performance of individual bats on each condition, and error bars indicate ±1 binomial standard deviation. Performance relative to baseline increased or decreased predictably based on the filtering of echoes. The pattern of results indicates that the upper and lower frequencies contribute to echo identification in separate, doubly dissociated ways. Grey dashed lines connecting columns indicate significance of exact binomial tests: ** = *P* < 0.001, * = *P* < 0.01.

## IV. DISCUSSION

### A. Amplitude discrimination performance is predictably affected by frequency filtering

As hypothesized, bats’ performance on the amplitude discrimination task on which they were trained (respond to S+, the stronger of two simultaneously-presented echoes; Fig. 1) was affected by the high- or lowpass filtering of the S+ and S- echoes. When S-, the weaker of the two echoes, was highpass filtered, the bats’ mean discrimination performance increased from 63% to 76%. Attenuating the lower frequencies of an incoming echo drastically disrupts the bat’s ability to perceive the precise delay (distance) of an ensonified target (Bates and Simmons, 2011; Ming *et al*., 2021). In the current experiment, attenuating just the bottom 10-15 kHz of S- resulted in an overall simpler S+/S- discrimination task for the bat and performance increased, despite the fact that the S- echo still contained a majority of its bandwidth and the same amount of acoustic energy as in the baseline condition. When the S- echo was instead lowpass filtered, mean discrimination performance also increased significantly, from 63% to 73%. Lowpass filtering of FM2 of incoming echoes, using a variety of filtering methods, has been shown to defocus the bat’s percept of echoes, making it difficult for them to determine the pulse-echo delay of filtered echoes and abolishing any masking effect the echo may have had before (Simmons et al., 2004; Stamper *et al*., 2009; Bates and Simmons, 2010; Bates *et al*., 2011). The same performance effect was seen here, as the bats were significantly less likely to perceive S- as the stronger of the two echoes when it was lowpass filtered (despite having the same amount of acoustic energy as when unfiltered). However, performance increased less in this condition than when S- was highpass filtered. This may be a result of the asymmetric perceptual roles of the bat’s harmonics: while filtering of the higher frequencies mimics off-axis clutter and results in defocusing of the bat’s sonar image, filtering the lower frequencies more drastically inhibits the bat’s perception of incoming echoes. This is also seen in the bats’ performance when S+, the higher amplitude echo, was highpass filtered. Mean performance decreased from 63% to 32%, showing that the bats actually reversed which echo they responded to, walking towards the weaker of the two echoes a majority of the time. The bats had a difficult time even perceiving the presence of the S+ echo, despite the fact that it remained an overall stronger acoustic signal than the S- echo. Anecdotally, the authors can report that all three bats that took part in that condition generally chose the S- echo quickly and with no hesitation, despite the continued lack of rewards for choosing that echo. These results once again highlight the disproportionate relevance of the lowest frequencies of the bat’s wideband FM calls to the bat’s biosonar perceptual system. Bates and Simmons (2010) found a similar reversal of discrimination performance when they highpass filtered their S+ echo at 66 kHz, such that it only contained FM2. In the current task, it took a much smaller amount of highpass filtering (set at 37-40 kHz, removing only the lowest ∼10-15 kHz) for the bats to reverse discrimination performance.

In the final permutation of filtering conditions, S+ was lowpass filtered at 70 kHz, resulting in a significant decrease in mean performance from 63% to 56%. The decrease in performance indicates that the bats had a harder time discriminating which echo was the stronger of the two. It can be assumed that the S+ echo was perceptually defocused as a result of its lowpass filtering, but this defocusing did not cause the bats to reverse performance to respond to S-, as when S+ was highpass filtered. Nonetheless, it became more difficult for the bats to decide which of the two echoes was stronger. One possible explanation for this is that the perception of an object’s size (partially a function of the reflected echo’s amplitude) also becomes defocused when lowpass filtered, along with the object’s perceived distance becoming blurred. This would result in small differences in object size being more difficult to discriminate, as seen in the S+ lowpass filtering condition. In this condition, the lowpass filtered S+ echoes were still recognized as echoes of comparable amplitude (indicated by the bats’ performance not falling below 50%), but the precise amplitude of the S+ echo may have been difficult for the bat to perceive, leading to lower discrimination performance.

It is of interest to note that the bats’ performance increased and decreased on the amplitude discrimination task following similar patterns as previous delay experiments, despite the fact that spectral filtering of the echoes presumably disrupts the bat’s percept of the echo’s delay, not its strength (i.e. its perceived size). This may be due to the fact that the echoes in the current task were presented at a constant delay throughout training and the experiment. Filtering of echoes affects the bats’ perception of that echo’s delay, which may make them less likely to respond to that echo regardless of the echo’s amplitude. For example, the bats’ increased performance in the S- highpass condition (relative to baseline) may be due to additive perceptual effects: not only is the S- echo slightly weaker than the S+ echo it has been trained to respond to, but now it also has a poorly-defined delay of *approximately* 4.84 ms, whereas S+ still has the well-defined delay of 4.84 ms, which it has had throughout the bat’s training. Thus, S+ is easier to choose as the correct echo because it is both stronger *and* has the precise delay the bat was trained to respond to. A possible control for this would be to train the bats to discriminate two echoes with different amplitudes but randomly variable delays, such that targets are presented within a certain range of pulse-echo delay values, which is changed randomly across trials or days. This would ensure that bats would learn to discriminate the two echoes based *only* on amplitude, without also inherently learning to respond to the specific pulse-echo delay values the echoes are presented at (as they may have in the current task). This would allow us to more precisely isolate the effects of filtering on amplitude discrimination without the possible confound of the bats learning to respond to specific pulse-echo delays (and thus not responding to echoes that do not have that precise pulse-echo delay). Unfortunately, COVID-19 restrictions made running these further controls not possible.

### B. Clutter rejection mechanisms are versatile across perceptual contexts

These results highlight the flexibility of the big brown bat’s clutter rejection mechanisms, which help them perceptually discriminate central target echoes from peripheral clutter echoes in the cluttered foraging scenarios among foliage that big brown bats are likely to encounter (Bates *et al*., 2011). When foraging among clutter, these bats will rapidly emit short, wideband FM calls during their pursuit and capture of small flying insects (Griffin, 1958; Neuweiler, 2000), within meters or centimeters of surrounding foliage. The big brown bat’s echolocation beam is about 110 degrees wide (−6 dB width; Hartley and Suthers, 1989), which results in the bat receiving a large number of echoes from all nearby surfaces to the front of the bat, not just the echoes from whatever surface it is directly aiming at. This constitutes the bat’s perceptual clutter problem: the bat must, quickly and accurately, perceive the precise location of an insect measuring no more than 1-3 cm in size based on the weak echoes the insect reflects, while also receiving a cascade of relatively stronger echoes created by surrounding foliage; these echoes may be offset from the insect’s location by any number of degrees, and may have higher, lower, or identical pulse-echo delays (from the bat’s perspective) as the insect echoes.

Importantly, not all frequencies of the bat’s FM calls are emitted with equal strength across the bat’s echolocation beam. The higher frequencies are emitted most strongly within the central 60 °, while the lower frequencies are emitted almost equally strongly across the entire echolocation beam (Hartley and Suthers, 1989; Bates *et al*., 2011). The result is that when a bat echolocates a nearby object located directly ahead of it, the target returns an echo that contains all of the frequencies in the bat’s original call. When a bat echolocates a nearby object that is located off-center, it instead returns an echo with attenuated higher frequencies relative to the low frequencies. This cue is very helpful for the bat, as psychophysical tasks have shown that attenuated FM2 frequencies result in an echo whose delay cannot be easily resolved (Bates *et al*., 2011). Thus, objects that are ensonified by the center of the bat’s beam (such as an insect being tracked), are well-resolved: the bat perceives its distance and location with a high degree of accuracy. Objects that are ensonified by the periphery of the bat’s beam (such as foliage to the side of the insect) reflect a lowpass filtered echo, which is still detectable but is defocused and more difficult to resolve its precise delay; in this way, these peripheral echoes do not mask the central object of interest, even if at similar pulse-echo delays and amplitudes.

Previous studies, using a variety of temporal discrimination tasks, have shown the efficacy of these clutter rejection mechanisms in defocusing lowpass filtered echoes so that the delay of unfiltered echoes can be perceived with a high degree of accuracy (Moss and Schnitzler, 1989; Simmons *et al*., 2004; Stamper *et al*., 2009; Bates and Simmons, 2010; Bates *et al*., 2011). Here, we extended these techniques of high- and lowpass echo filtering to an amplitude discrimination task and observed changes in discrimination performance that were in line with the conclusions from previous studies: lowpass filtering of a “clutter” or “distractor” echo (S-) led to the bats more successfully perceiving and choosing the “target” echo (S+), while highpass filtering either led to better discrimination performance (if S- was highpass filtered) or a reversal in performance (if S+ was highpass filtered). The current discrimination task the bats were trained on constituted a new perceptual context that has not been tested before, wherein both echoes from either side were presented simultaneously after every call, were spectrally identical, and had identical pulse-echo delay. The pattern of performance in the current task indicates that the bats’ perceptual clutter rejection mechanisms are adaptive not only when discriminating targets based on their delay (i.e. distance), but also when discriminating targets based on their amplitude (i.e. their size). Additionally, our results suggest that these clutter rejection mechanisms may not just modify their perception of echo distance, but also their perception of object size (as a function of echo amplitude).

## ACKNOWLEDGEMENTS

We thank Brandon Yeoh for assistance in bat training and data collection. This work was supported by an Office of Naval Research Multidisciplinary University Research Initiative grant (N00014-17-1-2736, J.A.S. and A.M.S.). A.T. is supported by a National Science Foundation Graduate Research Fellowship.

